# An Insectified Caco-2 Cell-Based assay to monitor pesticide Transport Processes and Pharmacokinetics

**DOI:** 10.64898/2026.07.21.739763

**Authors:** Kassiani Skouloudaki, Shane Denecke, Kathrin Vogelsang, Spyridon Pergantis, John Vontas

**Affiliations:** Institute of Molecular Biology and Biotechnology, Foundation for Research and Technology-Hellas, 71409 Heraklion, Greece; Department of Biology, University of Crete, 71409 Heraklion, Greece; Merck & Co., Inc. Lansdale, PA, 191446 United States USA; Bayer Crop Science, 40789 Monheim, Germany; Department of Chemistry, University of Crete, 71409 Heraklion, Crete; Department of Crop Science, Agricultural University of Athens, Athens, Greece

## Abstract

Intestinal insect cells are integral to pharmacokinetic research, unfortunately they do not serve as cell models for more advanced studies comparable to transport studies typically conducted in mammalian models. To overcome this limitation, in this study, we developed a Caco-2 cell line that has been genetically modified to mimic insect-like characteristics, providing a robust ‘insectified’ screening platform to monitor transport processes of xenobiotics. This platform consists of Wild Type (Pgpwild type) cells, an Pgp Knockout (PgpKO) line to eliminate endogenous background interference, and species-specific P-gp rescue lines. These rescue lines allow for the stable expression of human (Hs Pgp), cotton bollworm (Ha Pgp), and malaria mosquito (Ag Pgp) homologs, enabling a direct comparative analysis of efflux kinetics across different pesticide target and non-target biological systems. Functional validation using the substrate Digoxin confirmed high P-gp dependency within the platform. Methyl-parathion, an organophospate insecticide, was recognised and transported by both insect and human Pgps. This is consistent with the low mammalian selectivity of the compound. In contrast, Triflumezopyrim exhibited a high efflux ratio that remained remarkably stable even in the absence of MDR1, indicating that the uptake and pharmacokinetics of this compound are not Pgp-dependent. By successfully differentiating between transporter-specific and independent pathways these cell lines can serve as a high-fidelity screening tool for predicting novel pesticide selectivity and possibly validating the potential role of transporters in resistance. The system can be expanded and exploit also other transporters.

## Introduction

The intestinal epithelium and various physiological barriers serve as a primary barrier and determinant of the pharmacokinetics and toxicity of ingested environmental toxins, including a wide array of modern insecticides. While gut derived insect cell lines remain indispensable tools for pharmacokinetic screening, they possess structural limitations that impact their utility in complex physiological modelling. Most notably, these cells lack the organized epithelial architecture-characterized by polarized cell layers and robust tight junctions-that is essential for the barrier-function and transport studies typically conducted in mammalian systems. Without this specialized spatial orientation, insect cell cultures cannot fully replicate the selective permeability or the active transport mechanisms found in mammalian intestinal or blood-brain barriers. Because of these limitations, there is a need to develop better insect cell-like models.

One of the best *in vitro* models of intestinal epithelial cells available for studies of drug intestinal absorption and excretion is the human Caco-2 cell line [1–3]. In 1971, the Caco-2 cell line was established in culture from a human colon adenocarcinoma [4]. Caco-2 cells exhibit morphological as well as functional similarities to the human enterocytes [5]. When cultured under specific conditions, Caco-2 cells grow exponentially and, when in confluency, they undergo enterocytic differentiation, which is complete within 21 days in culture [4]. During their differentiation, they form a polarized monolayer that allow for the calculation of the efflux ratio (ER)-a key metric. This value allows to distinguish between active uptake, diffusion and active efflux. A value significantly greater than (>2) indicates active rejection by the cell. They also develop a well-defined brush border with regular microvilli on the apical surface, as well as tight cellular junctions [4,5].

Transporters have a major impact on the absorption, distribution, metabolism and excretion (ADME) of a diverse number of toxins. A critical component of this defense across species is the ABC transporters family. P-glycoprotein (P-gp: MDR1, ABCB1), an ATP-binding cassette (ABC)-transporter, is a membrane protein that utilizes ATP hydrolysis as the driving force for the efflux of drugs or xenobiotics from the cell, thereby limiting systemic absorption and preventing toxic accumulation within sensitive tissues [6]. P-gp is ubiquitously expressed on the apical surface of intestinal enterocytes [7] (Fig. 1A), the canalicular membrane of hepatocytes, the brush border membrane of renal proximal tubular cells and the endothelial cells of the blood brain barrier.

**Figure 1.**
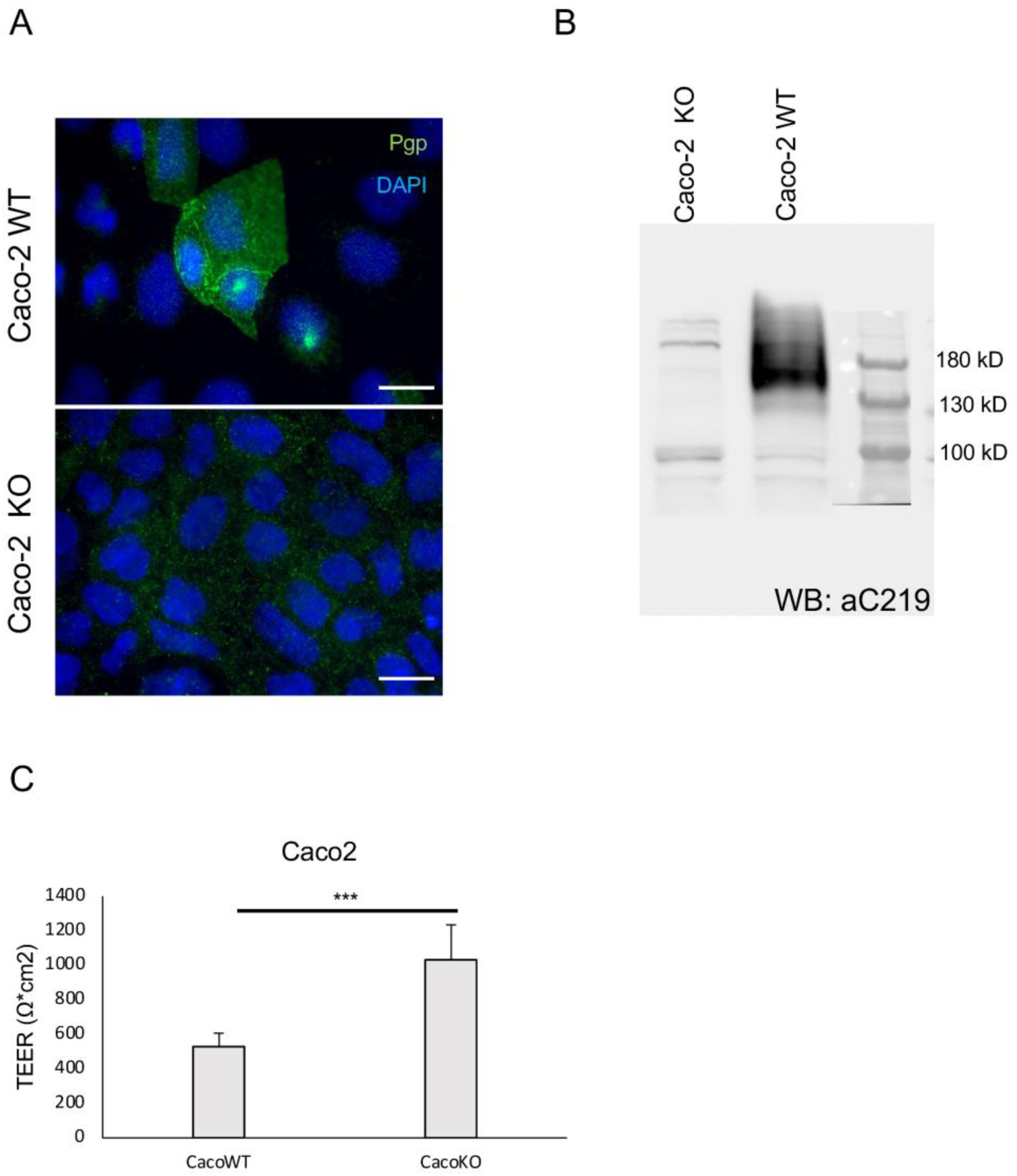
Validation of P-glycoprotein (P-gp) knockout in Caco-2 cells and its effect on monolayer integrity. A) Immunofluorescence microscopy of Pgp expression. Caco-2 wild-type (WT) and knockout (KO) cells were stained for P-gp antibody (green) and counterstained with DAPI (blue) to visualize nuclei. Scale bars 20μm. B) Western Blot (WB) analysis of P-gp protein levels. Cell lysates from Caco-2 KO and WT lines were probed with the C219 (Pgp) monoclonal antibody. C) Transepithelial Electrical Resistance (TEER) measurements. Comparison of monolayer barrier integrity between Caco-2 WT and KO cells. (***p < 0.001).

Membrane transporters, particularly the ABC (ATP-binding cassette) and SLC (Solute Carrier) families, serve as critical gatekeepers that mediate the pharmacokinetics of insecticide uptake across the insect gut and blood-brain barrier. As highlighted by Denecke et al [8] these transporters can actively efflux xenobiotics or limit their absorption, effectively reducing the internal bioavailability of the toxin before it reaches its molecular target. This ‘Phase III’ detoxification mechanism not only regulates the systemic distribution of insecticides but can also be a driver of pesticide resistance in several major agricultural pests. For example, the upregulation of Pgp homologs in the midgut and Malpighian tubules of pests such as the diamondback moth (*Plutella xylostella*) [9] or various mosquito species [10,11] is associated with increased tolerance to lethal doses of toxins like pyrethroids and avermectins. This evolutionary adaptation effectively lowers the intracellular concentration of the insecticide before it can reach its molecular target [12].

In the current study, we developed an “insectified” Caco-2 cell line, that can be used to study pesticide uptake selectivity across species-including target insect pests, humans, and non-target pollinators. This approach also serves to validate the role of certain Pgp genes (such as upregulated genes or different alleles) in conferring pesticide resistance.

## Results

### Validation of Caco-2 call lines and optimisation of permeability assays

To verify the successful depletion of Pgp transporter in the modified cell line its expression was confirmed using the C219 antibody through both immunofluorescence (Fig. 1A) and western blot (Fig. 1B) techniques. In the Caco-2 wild-type (Pgpwild type) line, human P-glycoprotein (Pgp/ABCB1) is clearly expressed at the cell membrane (Fig. 1A) whereas in the PgpKO cell line was found to be entirely absent, confirming the effectiveness of the knockout (Fig.1A). Western blot analysis showed a prominent band corresponding to P-gp in the Pgpwild type cell line between 130-180 kDa, representing the mature glycosylated form of the protein. This band is absent in the PgpKO cell line, confirming successful gene disruption (Fig. 1B). The integrity of the cell monolayer is measured by TEER. The PgpKO cells demonstrated higher TEER increases compared to wild-type cells (Fig.1C). Since a higher TEER value indicates a tighter, more effective barrier, this implied that better and faster tight junction formation occurred in those rapidly growing samples and that the knockout process did not cause deleterious structural defects.

#### Optimizing fluorescent compounds

We initially used the two commercially available cell lines, the Caco-2 wild-type (Pgpwild type) and the specialized P-glycoprotein (Pgp) depleted (PgpKO) cells. Rhodamin is a substrate for P-glycoprotein (Pgp) and can, therefore, be used as a molecular probe in compound-efflux assays. We first compared the efflux ratio of Rhod123 and RhodB along with the P-gp inhibitor, Verapamil in Pgpwild type line. The transport rate for the tested compound is expressed as the apparent permeability coefficient (Papp, Papp= dQ/dt *x* 1/A *x* C0). The efflux ratio (ER) was determined by the following equation ER= Papp (B-A)/ Papp (A-B), where Papp (B-A) is the Papp in the secretory direction and Papp (A-B) is the Papp in the absorptive direction.

Interestingly, we have observed that the efflux ratio of Rhod123 (ER >2) is significantly higher than the efflux ratio of RhodB in Caco-2 WT. Also, the presence of the Pgp inhibitor, Verapamil resulted in a notable decrease in the ER of Rhod123 whereas the ER of RhodB was only slightly reduced (Fig.2A). These results indicate that Rhod123 is the fluorescent compound to be used for further permeability assays.

**Figure 2.**
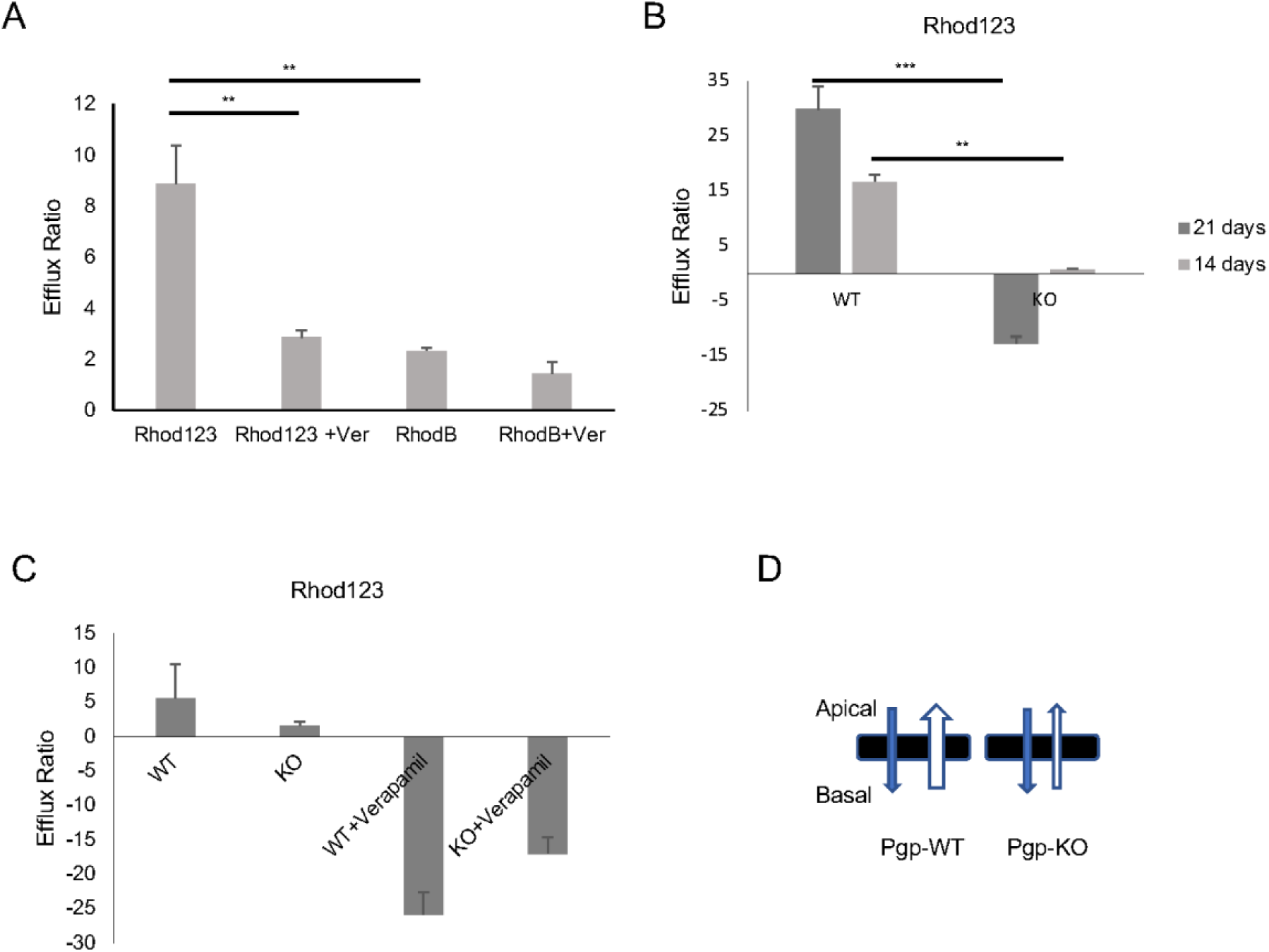
Effect of Pgp depletion on Rhodamine efflux or Functional activity of the P-glycoprotein (Pgp) efflux transporter using Rhodamine 123 (Rhod123) and Rhodamine B (Rhod B) as substrates. Α) Comparison of substrate specificity in Caco-2 wild-type cells using Rhod123 and RhodB. Β) Efflux ratios for Rhod123 in WT and KO cells measured at two different time points (14 and 21 days). C) Comparison of Rhod123 efflux in WT and KO cells with and without Verapamil. D) Diagram illustrating the directional movement of substrates across the apical and basal membranes in WT and KO cell lines.

#### Optimizing of seeding density

While Caco-2 cell monolayers remain stable for up to 30 days post-seeding, the traditional 21-day wait period restricts the frequency of testing. To optimize this timeline the most straightforward way to accelerate monolayer formation is to increase the initial cell number. Pgpwild type and PgpKO Caco-2 cells were cultured on transwells at 21,000 cells/ transwell for a 21-day growth period while a higher density of 40,000 cells/ transwell allowed for a shortened 14-day incubation. No significant difference was found between the tested cell densities. The Pgpwild type group shows high positive efflux ratios, meaning the transport system is functioning correctly and actively pumping Rhod123. Interestingly, the efflux activity increases significantly from day 14 to day 21, suggesting that the transport mechanism matures or becomes more active over time in the wild-type subjects. The PgpKO group shows a dramatic decrease in efflux. At 14 days, the ratio is near zero. By 21 days, the ratio becomes negative. This suggests that not only is the efflux pump missing, but there may be an accumulation of the dye within the cells or a complete reversal of the normal concentration gradient (Fig.2B)

#### Functional validation in Pgpwild type and PgpKO Caco-2 cell line

Caco-2 Pgpwild type and PgpKO cells were seeded as described and permeability assays were performed using Rhod123 as a fluorescent compound to validate the efflux ratio between Pgpwild type and PgpKO cell lines. To demonstrate proof of concept first the lucifer yellow assay and the TEER were used to measure the integrity of the monolayers for at least three independent experiments. Values ranging between 0.3-2% for lucifer yellow and > 300 ohms cm-2 indicate adequate monolayer integrity. All experimental values were within this range Next step was to perform compound-efflux assays using Rhod123 with and without P-gp inhibitor, Verapamil. We have observed significant changes in the efflux ratio of Rhod123 in Pgpwild type (ER >2) and PgpKO (ER < 2) cell lines (Fig. 2C) confirming the absence of Pgp in the knockout cell line. These data are demonstrated in the graphical model where in the Pgpwild type cells, the permeability from Basal-to-Apical (B to A) is much higher than Apical-to-Basal (A to B). In the PgpKO, these two values become nearly equal (Fig. 2D).

### Establishment and characterization of Caco-2 cells stably expressing *Helicoverpa armigera* Pgp (ABCB7), *Anopheles gambiae* Pgp (ABCBF) and *Homo sapiens* Pgp (ABCB1)

To establish the “insectified” screening platform PgpKO cells were transfected (Fig.3A) with either vector encoding full-length *Ha* Pgp, *Ag* Pgp or human *Hs* Pgp (as a positive control) with the proteins having a GFP tag (Fig.3B). Clones were selected based on G418 antibiotic selection and we chose one clone from each transgenic line with similar levels of expression for further study (Fig. 3A). Interestingly, the data presented in Figure 4 demonstrate that Pgp orthologs from humans and two insect species (mosquitoes and moths) can be successfully expressed in these cells. More specifically, immunofluorescent experiments using both C219 and GFP antibodies show Pgp (green) localized primarily at the cell membranes (Fig. 4A, C and E) whereas GFP reporter (green) localized ubiquitously throughout the cell (Fig. 4B, D and F) indicates successful clonal selection.

**Figure 3.**
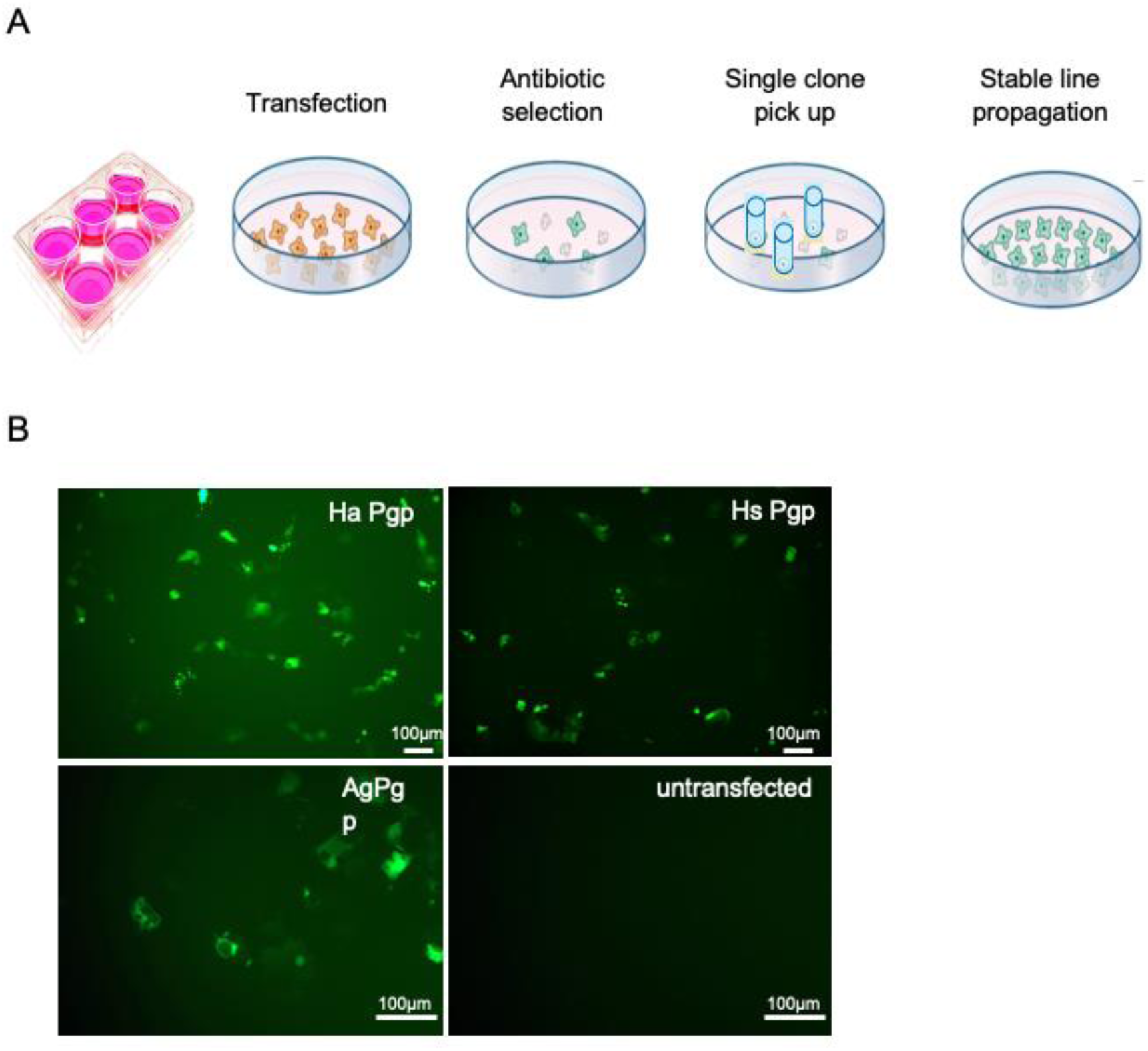
Establishment of engineered Caco-2 cell lines. A) Schematic illustration of stable cell line generation and single clone selection. B) Fluorescent images of positive clones. Scale bar: 100μm.

**Figure 4.**
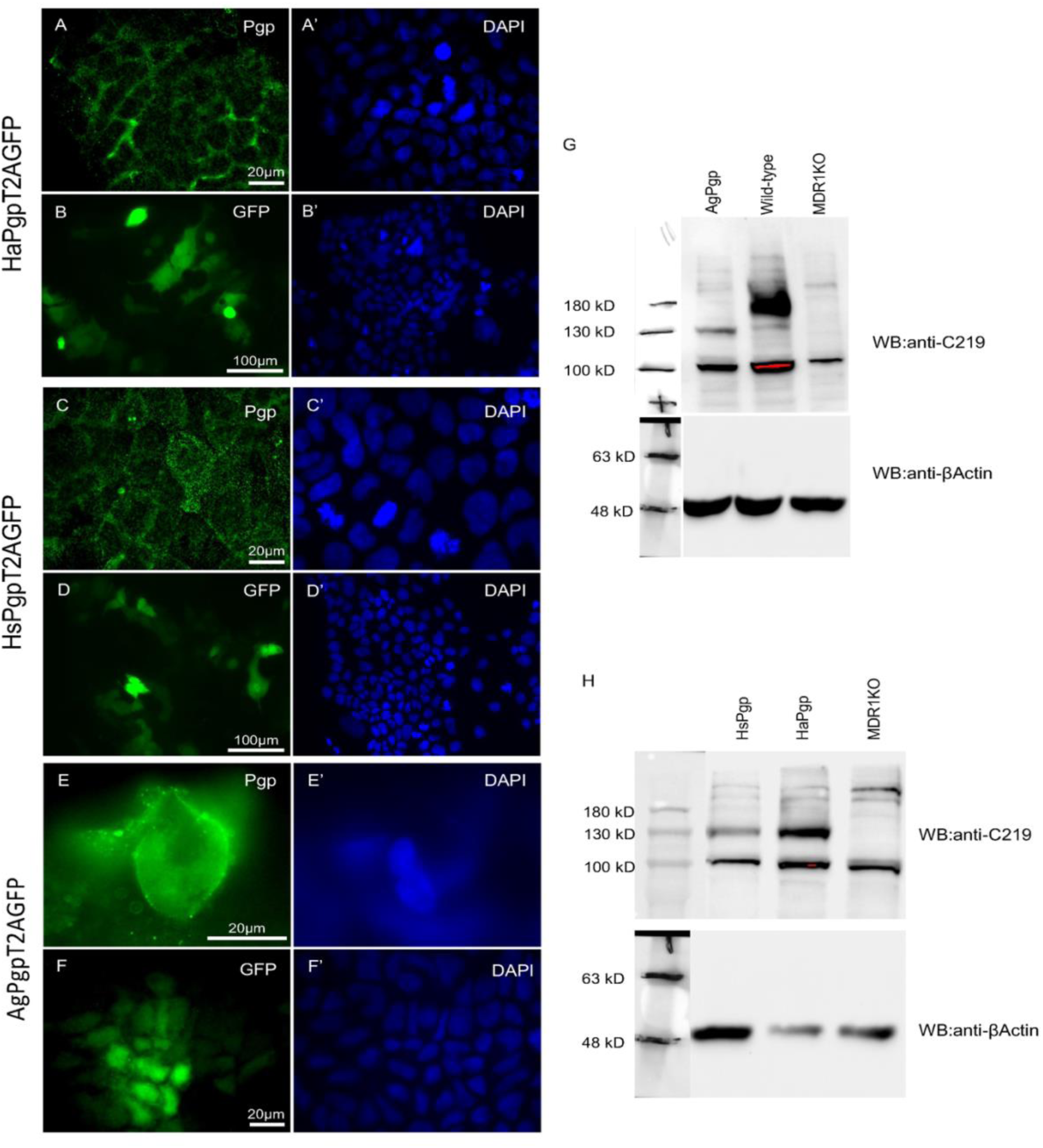
Analysis of P-gp expression and localization in engineered Caco-2 cell lines. Immunostaining of H. armigera (Ha. Pgp) expressing cells with antibodies against Pgp (A) and GFP (B). Nuclei were stained with DAPI (A’, B’). Immunostaining of H. sapiens (Hs. Pgp) expressing cells with antibodies against Pgp (C) and GFP (D). Nuclei were stained with DAPI (C’, D’). Immunostaining of A. gambiae (Ag. Pgp) expressing cells with antibodies against Pgp (E) and GFP (F). Nuclei were stained with DAPI (E’, F’). Scale bars: (A, A’, C, C’, E, E’, F, F’) 20 μm and (B, B’, D, D’)100 μm. Western blot from cell lysates showing the expression of different Pgp proteins from H. armigera, H. sapiens and A. gambiae in Caco-2 cells. Beta-actin is used as loading control (G, H).

Western blot analysis using the C219 antibody revealed significant bands between 130 kDa and 180 kDa, corresponding to the expected molecular weights of the Pgp orthologs (Fig. 4G, H). Specifically, the mature protein sizes are 139 kDa for HaPgp, 142 kDa for AgPgp, and 148 kDa for HsPgp, with HsPgp also visible in the wild-type (WT) lane next to AgPgp. Additionally, a distinct higher molecular weight band and smear is visible between 150–180 kDa, particularly in the wild-type lanes. This characteristic upward shift and smearing is a known result of N-linked glycosylation of P-glycoprotein.

### Enhanced Barrier Properties of Engineered Caco-2 lines for In Vitro Permeability Assay

The barrier integrity of the modified Caco-2 cell lines was evaluated through TEER measurements to ensure the formation of confluent, polarized monolayers. As illustrated in the Figure 5A, all engineered variants exhibited higher TEER values compared to the Pgpwild type (500 Ω*cm2), indicating that the genetic modifications did not compromise, and in fact enhanced, the tight junctional resistance of the monolayers. Notably, the PgpKO and HaPgp lines displayed a two-fold increase in resistance 1000-1150 Ω*cm2), suggesting a more restrictive paracellular pathway. While the human (HsPgp) and mosquito (AgPgp) orthologs showed more modest increases, all values exceeded the minimum threshold (typically >300 Ω*cm2) required for reliable drug transport assays. These results confirm that these cell lines provide a robust and viable physical barrier suitable for comparative permeability studies.

**Figure 5.**
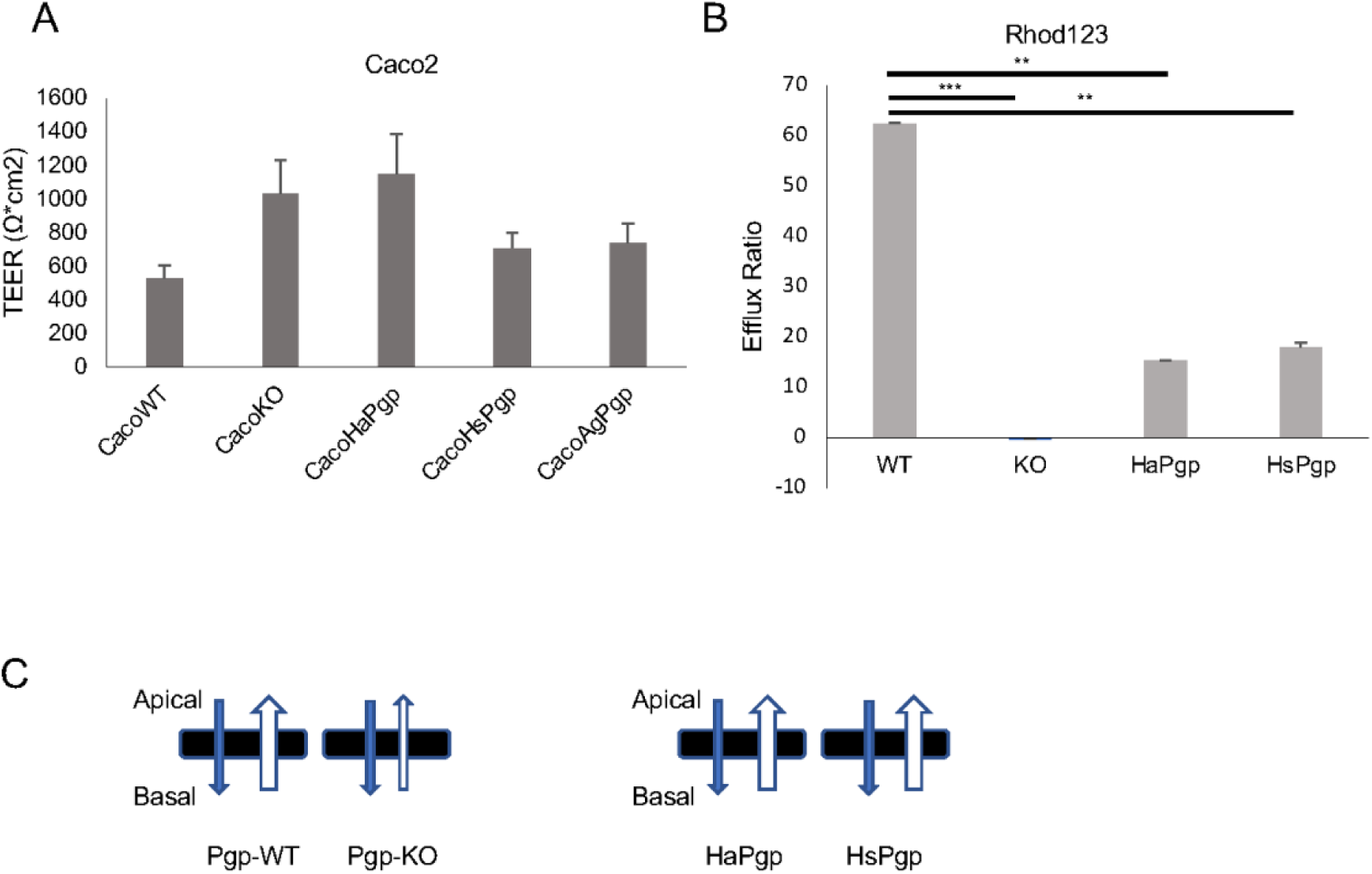
Barrier integrity and P-gp mediated efflux across the monolayers of engineered Caco-2 cell lines. A) Transepithelial Electrical Resistance (TEER) measurements. Comparison of monolayer barrier integrity between Caco-2 WT, KO and engineered cells. B) Comparative functional analysis of P-gp activity using Rhod123 as a substrate. (***p < 0.001, **p<0.01). C) Diagram illustrating the directional movement of substrates across the apical and basal membranes in WT, KO and engineered cell lines.

### *Helicoverpa armigera* Pgp orthologs can efflux Rhod123 as efficiently as the human Pgp

To evaluate the functional transport capacity of the P-glycoprotein (P-gp) variants, an efflux assay was conducted using Rhodamine 123 (Rhod123) as a fluorescent probe substrate. The Pgpwild type cells exhibited robust transport activity with a mean efflux ratio of approximately 63±0.5. In contrast, the PgpKO cell line showed a complete loss of transport function, with the efflux ratio reduced to negligible levels, confirming the necessity of the transporter for Rhod123 clearance. Heterologous expression of the moth (HaPgp) and human (HsPgp) orthologs successfully rescued the efflux phenotype, yielding ratios of 15.4 and 17.9, respectively (Fig. 5B). While both orthologs demonstrated significant transport activity compared to the PgpKO strain, neither reached the magnitude of efflux observed in the WT line, suggesting differences in protein expression levels or substrate affinity between the native and heterologous systems. These data are demonstrated in the graphical model where in the Pgpwild type cells, the permeability from Basal-to-Apical (B to A) is much higher than Apical-to-Basal (A to B). In the PgpKO, these two values become nearly equal (Fig. 2D).

### Comparative Analysis of Pesticide Efflux Activity

To investigate the role of P-glycoprotein (Pgp) in the membrane transport of different insecticidal compounds, we conducted comparative efflux assays using the various cell lines expressing Pgp transporters. We measured the efflux ratio (ER) of a classic substrate for P-glycoprotein (Pgp), Digoxin, a common insecticide, methyl-parathion and the mesoionic insecticide Triflumezopyrim. We observed that while the Pgpwild type cells maintain a high Digoxin efflux ratio of approximately 21, the complete removal of the transporter in the PgpKO cells brings this activity to nearly zero. The re-introduction of HaPgp, Ag Pgp and HsPgp into the cells successfully restored some transport function, resulting in efflux ratios of 4.2, 3.4 and 4, respectively (Fig. 6A). Methyl-parathion demonstrated a clear dependence on Pgp for cellular efflux, as evidenced by a significant reduction in efflux capacity upon Pgp deletion and subsequent functional rescue by human (*Hs*), moth (*Ha*), and mosquito (*Ag*) Pgp variants. Comparison of Pgpwild type vs. PgpKO showed that the ER dropped significantly from 4.67 in wild type cells to 2.52 in the PgpKO line. Expression of specific HaPgp, Ag Pgp and HsPgp homologues significantly restored efflux activity to 4, 4.24 and 4.41, respectively (Fig. 6B). Conversely, Triflumezopyrim exhibited negligible transport across the tested lines; no significant difference in efflux was observed between the Pgpwild type and PgpKO populations (p=0.13), and only marginal transport was recovered by the insect-derived *Ha* Pgp (Fig. 6C).

**Figure 6.**
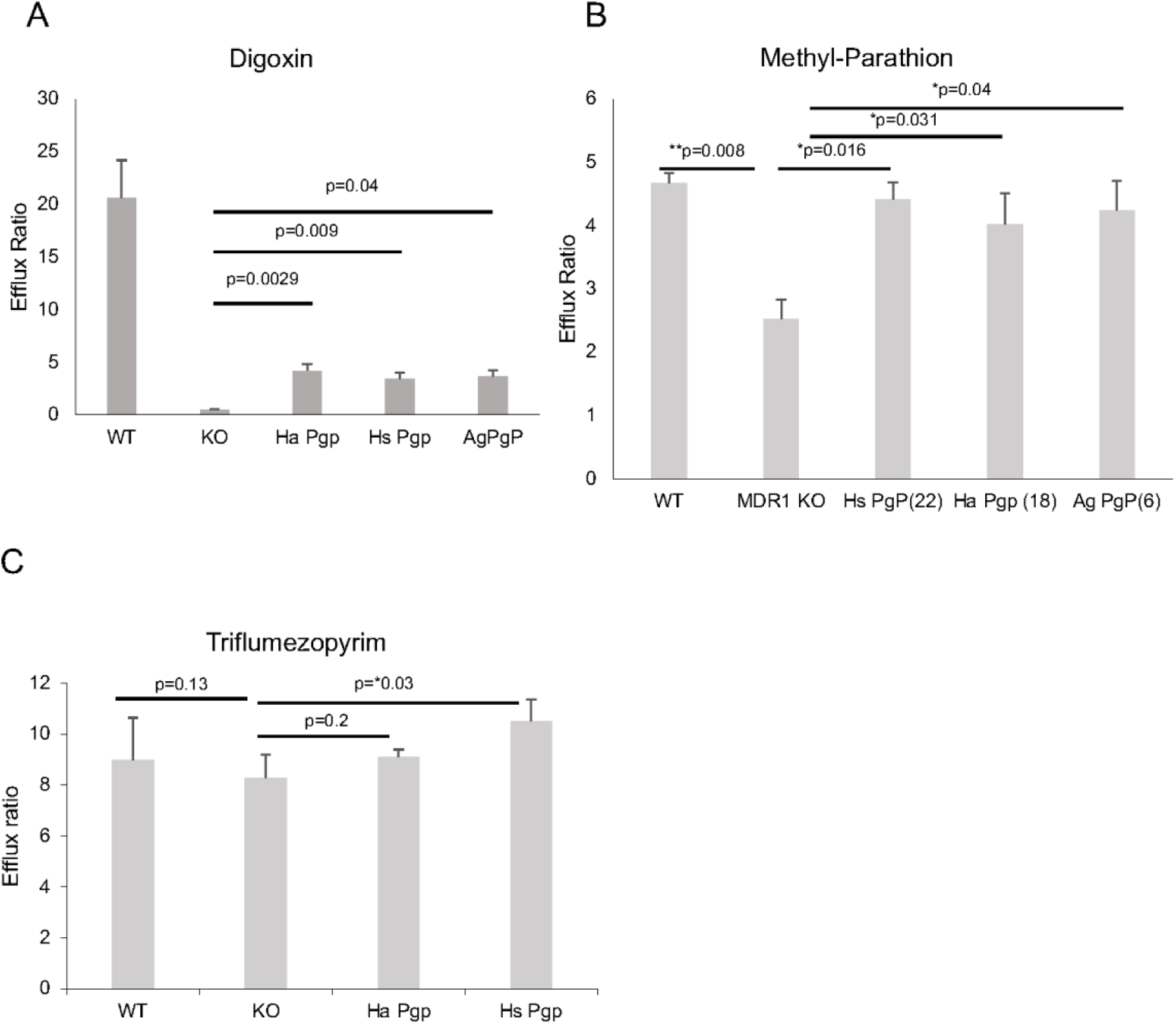
Assessment of H. armigera (Ha. Pgp), H. sapiens (Hs. Pgp) and A. gambiae (Ag. Pgp) Pgp-mediated transport. Efflux ratios of Digoxin (A), Methyl-Parathion (B) and Triflumezopyrim (C) substrates in WT, KO and engineered cell lines.

## Discussion

We successfully established and validated a robust in vitro permeability and efflux screening platform based on genetically engineered Caco-2 cells lacking endogenous P-glycoprotein (PgpKO) and stably expressing Pgp orthologs from humans (Homo sapiens), moths (Helicoverpa armigera), and mosquitoes (Anopheles gambiae). Initial characterization confirmed the effective knockout of Pgp in the Caco-2 PgpKO cell line. Immunofluorescence and western blot analyses demonstrated complete loss of Pgp expression in PgpKO cells, while wild-type Caco-2 cells showed strong membrane-localized Pgp expression, consistent with previous reports describing polarized ABCB1 localization in intestinal epithelial models [13]. Functionally, the absence of Pgp resulted in a near-complete loss of active efflux capacity, validating this line as a clean background for heterologous expression studies.

TEER measurements revealed that PgpKO cells exhibited higher TEER values compared to wild-type cells during monolayer formation. This observation suggests that deletion of Pgp does not compromise epithelial integrity; rather, it may enhance tight junction formation and barrier function. However, it is important to consider potential compensatory mechanisms, as the knockout of Pgp often triggers the upregulation of alternative efflux transporters (such as BCRP or MRPs) to maintain cellular homeostasis, Similar increases in TEER have been reported in other transporter-deficient epithelial models [14]. Importantly, all TEER values exceeded the threshold required for reliable permeability assays, ensuring that differences in compound transport were attributable to transcellular mechanisms rather than paracellular leakage.

Optimization of fluorescent probe selection demonstrated that Rhodamine 123 (Rhod123) is a superior Pgp substrate compared to Rhodamine B in Caco-2 wild-type cells. Rhod123 exhibited a markedly higher efflux ratio, which was strongly reduced in the presence of the Pgp inhibitor verapamil, confirming its specificity as a Pgp-dependent probe. In contrast, Rhodamine B showed only modest sensitivity to Pgp inhibition. These findings align with prior studies [15] and justify the use of Rhod123 as a sensitive reporter for subsequent functional assays. Functional validation experiments further confirmed the distinct transport phenotypes of Pgpwild type and PgpKO cells. While Pgpwild type cells exhibited high efflux ratios for Rhod123, PgpKO cells showed nearly equal apical-to-basal and basal-to-apical permeability, consistent with passive diffusion in the absence of active efflux.

A major advance of this work was the establishment of Caco-2 PgpKO cells stably expressing insect Pgp orthologs. Immunofluorescence and western blot analyses confirmed robust expression and correct membrane localization of H. armigera, A. gambiae, and human Pgp proteins. Importantly, TEER measurements showed that expression of these transporters did not disrupt epithelial barrier integrity; instead, all engineered lines exhibited equal or higher resistance compared to Pgpwild type cells, further supporting their suitability for permeability assays. Functional efflux assays revealed that H. armigera Pgp can transport Rhod123 with efficiency comparable to human Pgp, albeit at lower absolute efflux ratios than those observed in wild-type Caco-2 cells. The reduced magnitude of efflux in heterologously expressing lines likely reflects differences in expression levels, membrane density, or intrinsic substrate affinity between endogenous human Pgp and the introduced orthologs.

Digoxin, a canonical Pgp substrate, displayed high efflux ratios in Pgpwild type cells and almost complete loss of transport in PgpKO cells, validating the assay’s sensitivity. Partial restoration of Digoxin transport by insect and human Pgp orthologs confirms that these transporters retain the capacity to recognize and efflux structurally diverse substrates.

Methyl-parathion, an organophosphate insecticide, showed a clear dependence on Pgp for efflux. The significant reduction in efflux ratio upon Pgp deletion, followed by functional rescue with insect and human Pgp variants, suggests low selectivity. These data are in line with evidence that methyl parathion act as high-affinity inhibitors of P-glycoprotein (Pgp) [16–18].

In contrast, Triflumezopyrim exhibited minimal Pgp-dependent transport across all tested cell lines. The lack of significant differences between Pgpwild type and PgpKO cells indicates that this mesoionic insecticide is not a major Pgp substrate. However, Triflumezopyrim exhibited a high baseline efflux ratio of approximately 8 across all tested lines These results strongly indicate that Triflumezopyrim is a substrate for alternative active efflux transporters (such as BCRP or MRPs) rather than relying purely on passive diffusion or metabolic detoxification This distinction highlights the importance of compound-specific transport profiling when assessing resistance risk.

By combining a clean knockout background with heterologous expression of insect and human transporters, this system enables direct functional comparisons that are difficult to achieve in native tissues. Also, the significant reduction in efflux ratios upon Pgp deletion, followed by the functional rescue using insect Pgp variants could also possibly provide mechanistic insight into the role of efflux transporters in insecticide resistance. Over-expression of these transporters in insects could lead to a multidrug resistance, where the insect can effectively pump out various classes of insecticides before they reach their target sites.

Despite the versatility of the current Pgp transport assay, certain limitations exist regarding highly lipophilic, poorly water-soluble compounds. For such substances, non-specific binding to the plasticware or their retention by the cell monolayer can result in unreliable estimates of permeability. Similar challenges have been documented in Caco-2 cell-based systems, where the addition of bovine serum albumin (BSA) to the receiver compartment was utilized to maintain sink conditions [19]. Implementing these steps in our current model could ensure to distinguish Pgp-mediated transport from passive membrane retention.

## Materials and methods

### Cell lines

Caco-2 (CRL2102) was purchased from ATCC and Caco-2 MDR1 KO (MTOX1001) was purchased from Sigma-Aldrich (St. Louis, MO). CRL2102 cells were grown in DMEM 11995 supplemented with 10% fetal bovine serum (FBS) and 0.01mg/mL apo-transferrin (cat no T539). MTOX1001 were cultured in DMEM D5671 supplemented with 20% FBS (only for the first two passages, cat no F4135) and then 10% FBS, 2mM Glutamine (cat no G7513) and gentamycin.

### Chemicals

Chemicals and reagents were purchased from the following providers: Rhod 123 (Cat. No. 83702), Rhod B (Cat. No. R6626), Verapamil (Cat. No. v4629), Lucifer yellow (Cat. No. L0259) and Glucose (Cat. No. G7021) Sigma-Aldrich (St. Louis, MO, USA); HBSS: Gibco, Thermo Fisher Scientific Waltham, MA, USA; Cat. No. 14025-050; Methyl Parathion-d6: LGC Standards, Teddington, UK; Triflumezopyrim: LGC Standards, Teddington, UK; Digoxin: Sigma-Aldrich, St. Louis, MO, USA; Cat. No. D6003.

### Transient transfection and generation of stable cell line

The coding sequence of Helicoverpa armigera (HaOG200306) ABCB7, Anopheles gambiae (AGAP005639) ABCBF and Homo sapiens ABCB1 were synthesized and cloned in pcDNA3.1 vector (Genscript, Piscataway, NJ). Plasmids were linearized with a single cutter enzyme and purified with Phenol-chloroform. Afterwards cells were plated in a 6-well plate for transfection. Next day 3μg of linear *empty pcDNA3.1 vector or with vector containing full-length ABCB7, ABCBF or ABCB1* per well were used along with the Transfex ATCC transfection reagent to transfect the Caco-2 Pgp KO (MTOX1001) line.

The medium was changed 24 hours post transfection and cells were left to grow for a total of 72hours. Cells were transferred from the 6 well plate into 150mm plates. Cells from one 6 well were transferred in two 150mm plates. At the same time, we started selection with selection antibiotic using the concentration we have chosen for the specific cell line after titration experiment (G418, 500ug/mL). For clonal selection medium was changed every 3 days. An untransfected cell culture control was transferred from 6-well to 10cm plate and subjected to the same antibiotic regime. The control was used to confirm the efficacy of selection, as no colonies were formed under these conditions. The selection period lasted for approx. 20 days. First clones started to be visible and ready to picked up at around 14-15 days. Single clones were transferred into 12 well plates. As soon as a clone in the 12 well plate became confluent it was transferred it to a 25cm2 flask. When the 25cm2 flask became confluent cells were frozen in two aliquots and a small part was kept for protein extraction.

### Protein extraction and Western blot

Cells were lysed in 10 mM NaCl, 25 mM HEPES pH 7.5, 2 mM EDTA and a mammalian protease inhibitor cocktail. After centrifugation, membrane proteins were solubilized in 100 mM NaCl, 25 mM HEPES pH 7.5, mPIC and 1% (w/v) (3-((3-cholamidopropyl)-dimethylammonio)-1-propanesulfonate, (CHAPS)). Boil at 65°C for 10min using loading buffer with mercaptoethanol. Proteins (30 mg per lane) were separated by sodium dodecyl sulfate polyacrylamide gel electrophoresis (SDS-PAGE; 10% gel) and transferred onto nitrocellulose membrane 0.2um pore size, wet transfer 2.5h at 250mA. Non-specific binding sites were blocked by incubation for 60min in 4% (w/v) dry milk and 1% BSA in TBS; 150 mM NaCl, 50mM Tris pH 7.5) at room temperature. The membrane was incubated with primary antibodies a-Pgp C219 (Thermo Fisher Scientific, Waltham, MA, USA, Cat. No. MA1-26528) in dilution 1:200 and beta-actin D6A8 (Cell Signaling Technology, Danvers, MA, USA; Cat. No. 8457) in dilution 1:1000 in TBS containing 0.3% (v/v) Tween-20 overnight at 4° C. After rinsing the membrane three times with TBST for 15 min each, secondary antibodies conjugated to horseradish peroxidase were incubated for 1 hour at room temperature. After rinsing the membrane three times with TBST for 10 min each, signals were visualized with an enhanced chemiluminescence detection system (Supersignal West femto, Thermo Fisher Scientific, Waltham, MA, USA)

### Immunofluorescence

Caco-2 cells were fixed in a 4% PFA for 20 min, permeabilized with 0.2% Triton X-100, and blocked in PBS containing 0.2% horse serum. Immunostainings for Pgp and GFP were performed with a mouse monoclonal P-glycoprotein antibody, MA5-13854, Invitrogen) at a dilution of 1:50 and a rabbit polyclonal a-GFP antiserum 4B10 (Cell Signaling Technology, Danvers, MA, USA; Cat. No. 2955) at a dilution of 1:50, in combination with fluorescently labelled secondary antibodies. Images were obtained with an LSM 510 confocal microscope (Zeiss).

### Lucifer yellow (LY) quantification

*Lucifer yellow (*LY) quantification was performed by the addition of 75 μL of 60 μM LY solution (in HBSS with 1% DMSO) to the apical side of each well. Incubate the plate at 37°C, 5% CO2 for 1 hour while shaking (40-50 rpm). During incubation, LY 1:3 serial dilutions were made for standard curves: starting with 60 μM up to 6.7 μM as a high standard curve. For a low standard curve 1 μM was used as a starting concentration and a series of 1:3 dilutions were followed up to 4.1 nM. Three wells of HBSS 1% DMSO were included as a blank.

Aliquots from the receiver wells (100 μL) and transwell inserts (50 μL) were transferred to solid black plates; the latter were diluted 1:1 in HBSS with 1% DMSO. Fluorescence intensity was measured via a plate reader using an excitation/emission filter set of 480/530 nm. Then the following calculations were made:

% LY rejection = 100 [1 – RFUbasolateral / RFUapical]

### TEER assay

The integrity of the monolayers was monitored by measuring transepithelial electrical resistance (TEER) before the bidirectional transport experiments using a EVOM3 with STX2-Plus electrode epithelial voltmeter (World precision Instruments-WPI, United States). The transporter experiment was only conducted on cell monolayers with TEER values above 300 Ω*cm2 [20].

### Surface treatment for transwell

Transwell membranes were coated with Collagen I (Ibidi, Gräfelfing, Germany) at a surface concentration of 5 μg/cm². The Collagen I stock was diluted in 17.5 mM acetic acid to prevent premature polymerization. Each transwell received 100 μL of the collagen working solution and was incubated for one hour at room temperature. Following incubation, the solution was removed, and the membranes were washed three times with DPBS. Coated transwells were either used immediately or air-dried overnight at 4°C. Prior to cell seeding, the treated membranes were equilibrated with cell-specific media for a minimum of 15 minutes.

### Transwell assay and bidirectional transport

#### Assay conditions (APICAL TO BASAL) or (BASAL TO APICAL)

Permeability assays were conducted in triplicate using HBSS buffer supplemented with 10 mM D-glucose and 20 mM HEPES. Test compounds, including Rhodamine 123, Rhodamine B (30μM), Digoxin, Triflumezopyrim, and Methyl-parathion (10μM), were prepared in DMSO. At t =300μL of the compound working solution was added to the apical compartment and 800μL of buffer to the basal compartment, after which 50μL was immediately recovered from each for initial concentration verification.

After a 2-hour incubation, samples were collected from both compartments for analysis. To prepare for quantification, 100μL from the receiver wells and 50μL from the donor inserts (diluted 1:1 in HBSS with 1% DMSO) were transferred to solid black plates. Αbsorption was measured using a SpectraMax M2e photometer at specific wavelength pairs: 498/530 nm (Rhodamine 123), 546/568 nm (Rhodamine B), and 480/530 nm (Lucifer Yellow).

#### Transwell data analysis

*P*app values of the test compounds were determined using the following equation: (Papp=ΔQ/Δt *x* 1/A *x* C0), where ΔQ / \Δt represents the rate of drug transport across the membrane over the 2-hour incubation period. A is the surface area of the transwell insert (0.33cm2 for standard 24-well inserts). C0 is the initial donor concentration in the donor chamber at t = 0.

#### LC/MS

##### Quantitative Determination of Parathion-methyl-d6

The quantitative determination of Parathion-methyl-d6 was conducted using reversed phase high performance liquid chromatography (HPLC) with high resolution Orbitrap mass spectrometry. More specifically, a microbore C18 reverse phase column (2.1 mm × 10 mm, 3 μm particle diameter, Fortis) was used with a mobile phase composed of H2O with 0.1% formic acid (FA) (Solvent A), and acetonitrile with 0.1% FA (Solvent B). The gradient composition was as follows: 0 min to 7 min solvent A decreased from 95% to 20%, whereas solvent B increased from 5% to 80%, this composition remained constant for another 2 min, i.e. until min 9. From 9 to 10 min solvent A increased from 20 to 95%, whereas solvent B decreased from 80% to 5%. This composition remained constant for another 3 min prior to the next injection. Mobile phase flow rate was at 200 μL/min, and samples were injected at a volume of 2 μL.

Analyte detection was carried out using a QExactive Plus Orbitrap mass spectrometer operating in its targeted single ion monitoring (tSIM) mode at the mass of protonated Parathion-methyl-d6, which has an elemental composition of [C8H4D6NO5P + H]+ and a m/z of 270.0467 ± 0.0015. Adequate mass accuracy was achieved at an Orbitrap resolving power of 140,000. Mass spectrometer mass calibration was conducted daily. Standards were prepared at concentrations ranging from 0.1 to 1.0 μm Parathion-methyl-d6 and all samples were diluted accordingly to have their concentrations fall within this concentration range.

##### Quantitative Determination of Digoxin

The quantitative determination of Digoxin was conducted using reversed phase HPLC with high resolution Orbitrap mass spectrometry, using a microbore C18 reverse phase column (2.1 mm × 10 mm, 3 μm particle diameter, Fortis) with a mobile phase consisting of H2O with 0.1% formic acid (FA) (Solvent A), and acetonitrile with 0.1% FA (Solvent B). The gradient composition was as follows: 0 min to 1 min solvent A remained at 90%, whereas solvent B at 10%. From 1 to 3 min solvent A decreased to 10%, whereas solvent B increased from 10% to 90%, this composition remained constant for another 4 min, i.e. until min 7. From 7 to 8 min solvent A increased from 10 to 90%, whereas solvent B decreased from 90% to 10%. This composition remained constant for another 3 min until the next injection. Mobile phase flow rate was at 200 μL/min, and samples were injected at a volume of 2 μL.

Digoxin detection was carried out using the QExactive Plus Orbitrap mass spectrometer operating in full MS mode, scanning from 770 to 820 m/z. Digoxin, which has an elemental composition of C41H64O14, was detected as [C41H64O14 + Na]+ with m/z of 803.4188 ± 0.0025. Adequate mass accuracy was achieved operating at an Orbitrap resolving power of 140,000. Mass spectrometer mass calibration was conducted daily during this analysis.

Standards were prepared at concentrations ranging from 0.1 to 1.0 μm Digoxin and all samples were diluted accordingly in order to have their concentrations fall within this linear response range.

##### Quantitative Determination of Triflumezopyrim

The quantitative determination of Triflumezopyrim was conducted using reversed phase HPLC with high resolution Orbitrap mass spectrometry. More specifically, a microbore C18 reverse phase column (2.1 mm × 10 mm, 3 μm particle diameter, Fortis) was used with a mobile phase composed of H2O with 0.1% formic acid (FA) (Solvent A), and acetonitrile with 0.1% FA (Solvent B). The gradient composition was as follows: 0 min to 1 min solvent A remained at 90%, whereas solvent B at 10%. From 1 to 3 min solvent A decreased to 10%, whereas solvent B increased from 10% to 90%, this composition remained constant for another 4 min, i.e. until min 7. From 7 to 8 min solvent A increased from 10 to 90%, whereas solvent B decreased from 90% to 10%. This composition remained constant for another 3 min until the next injection. Mobile phase flow rate was at 200 μL/min, and samples were injected at a volume of 2 μL.

Triflumezopyrim (C20H13F14N4O2) detection was carried out using the QExactive Plus Orbitrap mass spectrometer operated in parallel reaction monitoring (PRM) mode with m/z 399.11 being the precursor ion with a 1 m/z isolation window. Fragmentation was carried out in the higher-energy collision dissociation (HCD) at a ce of 27. The reaction product ions were 279.0737 ± 0.0030 and 121.0398 ± 0.0030. Adequate mass accuracy was achieved operating at an Orbitrap resolving power of 140,000. Mass spectrometer mass calibration was conducted daily.

Standards were prepared at concentrations ranging from 0.1 to 1.0 μm Triflumezopyrim and all samples were diluted accordingly in order to have their concentrations fall within this linear response range.

## Acknowledgments

We are grateful to the Bayer team members Panagiota Papazoglou and Jinal Badrakia for their technical support. The project was funded by BAYER AG (agricultural pest transporter work). Additionally part of this work was supported by the Gates Foundation INV-079677. The conclusions and opinions expressed in this work are those of the author(s) alone and shall not be attributed to the Foundation.

